# A thermodynamic model of bacterial transcription

**DOI:** 10.1101/2021.10.27.466199

**Authors:** Jin Qian, David Dunlap, Laura Finzi

## Abstract

Transcriptional pausing is highly regulated by the template DNA and nascent transcript sequences. Here, we propose a thermodynamic model of transcriptional pausing, based on the thermal energy of transcription bubbles and nascent RNA structures, to describe the kinetics of the reaction pathways between active translocation, intermediate, backtracked, and hairpin-stabilized pauses. The model readily predicts experimentally detected pauses in high-resolution optical tweezers measurements of transcription. Unlike other models, it also predicts the effect of tension and the GreA transcription factor on pausing.

## I. INTRODUCTION

During bacterial transcription, there are frequent pauses in the forward translocation of RNA polymerase. Pauses have been observed widely *in vivo* and *in vitro* and vary in durations from milliseconds to minutes [1, 2]. Long pauses, which may last tens of seconds, are classified as Class I ‘hairpin-stabilized’ and Class II ‘backtracked’ signals. Short pauses, which typically last less than one second, are proposed to be intermediate precursors of long pauses [3]. Class I and Class II pauses have been structurally characterized and mechanistically explored [4, 5]. They are thought to be regulated by the sequence of the DNA template, the structure of the nascent transcript, and the availability of transcription factors [6–8].

Previous models of the kinetics of backtracked pauses reproduced some types of experimentally detected pauses [9–12] but fail to predict other types of pausing and pause duration, and do not treat external tension or transcription factors. Here, we propose a model based on our current biochemical understanding of transcription pausing mechanisms and optimize the parameters of the model with high-resolution transcription data. After training, this purely thermodynamic model accurately reproduces experimentally observed pause sites and durations, and provides a mechanistic explanation of the effect of external tension and transcription factors. Furthermore, the model accurately predicts transcription dynamics on random DNA sequences and is readily extendable to incorporate the initiation and termination stages.

## II. MODEL DESCRIPTION

The energy of the ternary elongation complex (TEC) is estimated as the sum of four contributions: the free energy of the (i) transcription bubble, (ii) DNA-nascent RNA hybrid, (iii) nascent RNA, and (iv) RNAP-DNA :

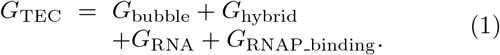

In this estimate, the first two terms are clearly sequence-dependent, and the secondary structure of nascent RNA (the third term) is also directed by sequence. The fourth term corresponds to interactions between the nucleic acids and RNAP subunits and is considered sequence-independent (effectively constant, as in Yager&von Hippel, Tadigotla, and Bai) [9–11]. A TEC is described by a transcription position (*m*) and state (*n*). The position along the template (*m*) indicates the length of the RNA transcript. TEC can be in either one of two translocation states: active (*n*=0) and backtracked (*n*<0), or in a conformational state, the hairpin-stabilized State (*hs*p). The interconnection among these states is shown in Figure 1a. From an active state at position *m* (*m*,0), a transcription complex can enter the next active state (*m*+1,0), or branch into backtracked state (*m*, −1) or hairpin-stabilized state (*m, hsp*).

**FIG. 1.**
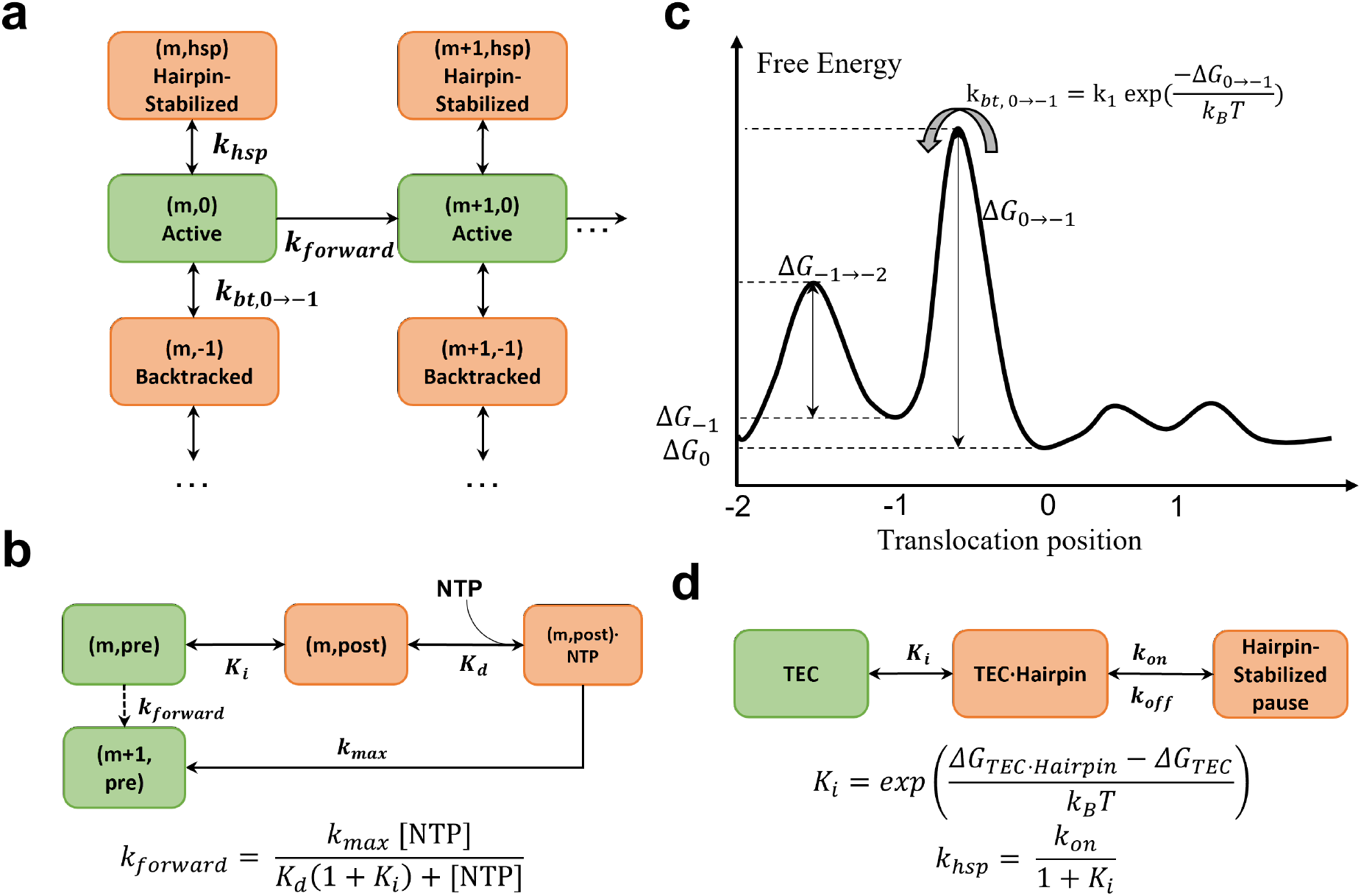
Model description: (a) A diagram of transcriptional states considered in the model and their interconnections. (b) An illustration of the RNAP forward translocation using Michaelis equation. (c) The free energy landscape for the backtracking pathways. Note that the first backtracking step has different energy barrier than the deeper backtracking steps. (d) The proposed kinetic mechanism for the hairpin-stabilized pause and its kinetics.

To determine the configuration of a transcription bubble and the details of the energy profile of a TEC, we used an approach based on statistical mechanics, the basis of which was described by Tadigotla [10]. The details are given in the SI.

Briefly, the active translocation of RNAP is modeled by the Michaelis-Menten (M-M) equation

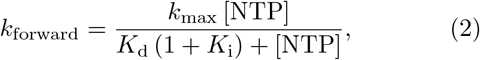

where *k*_max_ is the rate of NTP catalysis, *K*_d_ is the NTP dissociation constant, and *K*_i_ is the equilibrium constant between two adjacent translocation states (Figure 1b). This is a good approach, whether the translocational register of a TEC is considered to be determined by the presence of incoming NTP, as in the Brownian-ratchet model [13], or by the release of pyrophosphate [14], as long as *k*_max_ and *K*_d_ are given a different physical meaning in each case (SI).

Backtracking has been previously modeled using the Arrhenius Equation (3) with an activation barrier of 40 − 50 *k*_B_*T* for each step of backward translocation [9]. This value seems unreasonably high given that the free energy of base pairing in a transcription bubble is typically less than −20 *k*_B_*T* [10].

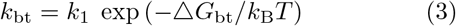

We take the same Arrhenius approach but treat the first step of backtracking differently from the subsequent ones (Figure 1c), based on the assumption that initially the 3’ end of the nascent transcript disrupts the active site and invades the secondary channel of RNAP [15], while additional backtracking stabilizes the interaction of RNA within the secondary channel. The model estimates an energy barrier about 10 k_B_T for the first step of backtracking and 2 k_B_T for the subsequent steps. The details are given in the SI.

Hypertranslocation refers to the forward translocation of RNAP without concurrent RNA elongation at the active site. However, we do not consider hypertranslocation for two reasons. First, hypertranslocation may not be a general phenomenon during transcription [16], and it cannot be distinguished from backtracking in force spectroscopy assays. Second, hypertranslocation is never energetically favored, because the number of base-pairing is reduced with respect to the active state.

To model a hairpin-stabilized pause, we take an allosteric view, in which an RNA hairpin contacts a short *α* helix at the tip of the RNAP flap domain that covers the RNA exit channel to induce the pause [4, 17]. The pathway is modeled as a fast equilibrium between two configurational states, a state free of hairpin and a state with a hairpin positioned close to the RNAP flap domain. The equilibrium is followed by a rate-limiting catalytic step (Figure 1d). The equilibrium is considered rapid compared to the formation of chemical bonds that stabilize the inactive state (see SI for details).

Since the experimental data we used to validate the model were acquired under tension of magnitude up to 25pN, the effect of external tension on the thermodynamics of TEC was considered. For the forward translocation and backtracking pathways, we employed the idea that the energy barrier is modulated by the work produced by tension [18]. The hairpin-stabilized pause was assumed to be unaffected by any applied tension, since it does not involve RNAP translocation.

It is important to notice that transcription is a process that involves only very small numbers of reacting molecules, thus the law of mass chemical reaction is not suitable. Rather, we apply two stochastic kinetic methods: (i) the continuous-time Markov chain and (ii) stochastic simulation. The continuous-time Markov chain allows us to analytically solve for the expected time spent in each state at a certain position (SI). The stochastic simulation reveals how individual pausing events develop.

The model is encapsulated in a MATLAB class object, which can generate a predicted residence time histogram with the input of a template sequence and a guess of unknown parameters (Table S2). Thus, the model can be trained with the data from real-time single molecule experiments. We used the time series obtained in high-resolution optical tweezers transcription experiments by Gabizon et al. [19] with or without GreB and RNase A that are known to interact with RNA and affect RNAP pausing. The transcription experiments were performed on a DNA template (8XHis) containing the T7A1 promoter followed by eight tandem repeats of a 239 bp sequence containing the *his*-leader pause site and four other known sequence-dependent pause sites [1]. The temporal resolution is high enough to detect pausing events longer than 100 ms. This allows sampling of the residence time with one base-pair resolution. Alignment of the traces under different forces and with different transcription factors generates the residence time histograms (Figure 2) as described previously.

**FIG. 2.**
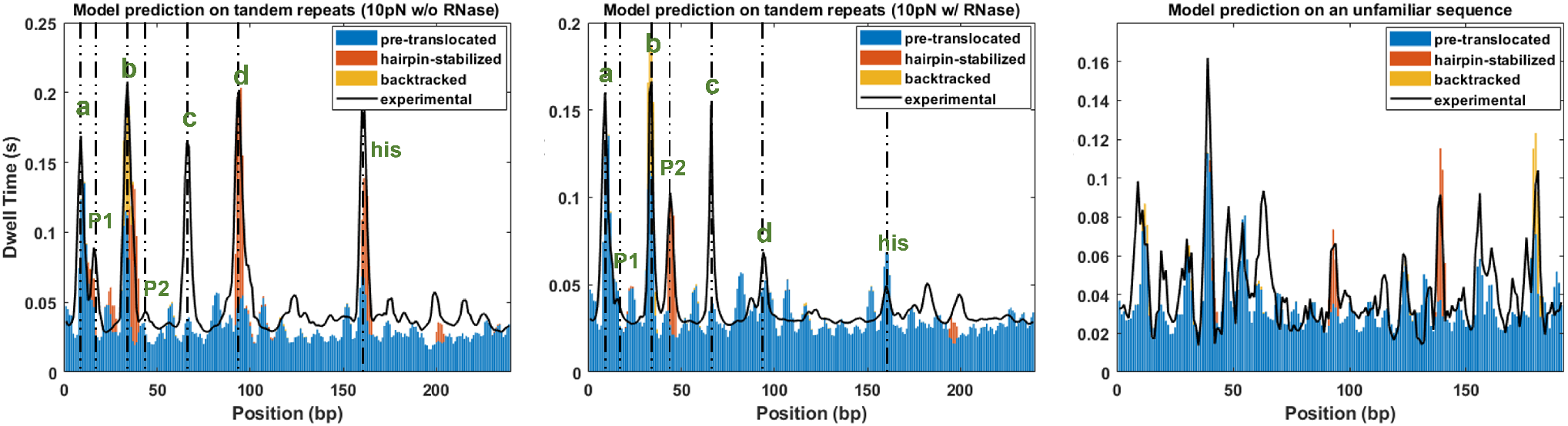
Model fitting and prediction. (a) Stacked histogram produced by the model for the condition of 10 pN in presence of RNase. The residence time due to different pausing mechanisms is represented by different colors. The experimental result is shown by the black line. (b) Stacked histogram produced by the model for the condition of 10 pN in absence of RNase. (c) Predicted histogram by the model on an unfamiliar sequence.

## III. COMPARISON OF THE MODEL WITH EXPERIMENTAL DATA

Experimental data under different conditions with various accessory factors helped to manifest the mechanisms of the pauses. Also, the analysis of the backtracking dynamics helped differentiate backtracked pauses from others. Table S1 summarizes the position and duration of pauses as well as their response to GreB or RNase (factors). Pauses at position ‘a’ are likely pre-translocation, since their duration is barely affected by the addition of GreB or RNase. Pauses at position ‘b’ are likely due to both backtracking and hairpin-stabilization, as their duration responds to the presence of GreB and RNase, and they are preceded by backward RNAP translocation, as previous analysis suggests [19]. ‘P1’, ‘d’ and ‘*his*’ are hairpin-stabilized pauses that almost disappear in the presence of RNase. Pause ‘P2’ is also hairpin-related, but unlike hairpin-stabilized pause ‘P1’, ‘d’ and ‘*his*’, it only appears in the presence of RNase.

We optimized the values of the model parameters to yield a residence time histogram that resembled the experimental data (Figure 2a and b). Since the model includes many parameters, we avoided overfitting by decomposing the overall kinetics into three parts that correspond to the pausing mechanisms and sequentially tuning the parameters related to each pausing mechanism (SI). The model clearly reproduces the positions and lifetimes of pauses observed experimentally except for pause ‘c’.

The model also successfully predicts the mechanisms of the pauses suggested by the experimental results (Figure 3). The experimental data suggest that the presence of GreB extends the dwell time at pause ‘b’ [19]. The model achieves this effect by adjusting the energy barrier of backtracking. The presence of RNase significantly decreases the dwell time at sites ‘P1’, ‘d’ and ‘*his*’, increases the dwell time at sites ‘P2’, and has little effect on the duration of pauses at other sites. RNase effectively shortens the lengths of nascent RNA available to form secondary structure, and constraining the model to operate on shorter nascent RNA reproduces the observed changes in pauses.

**FIG. 3.**
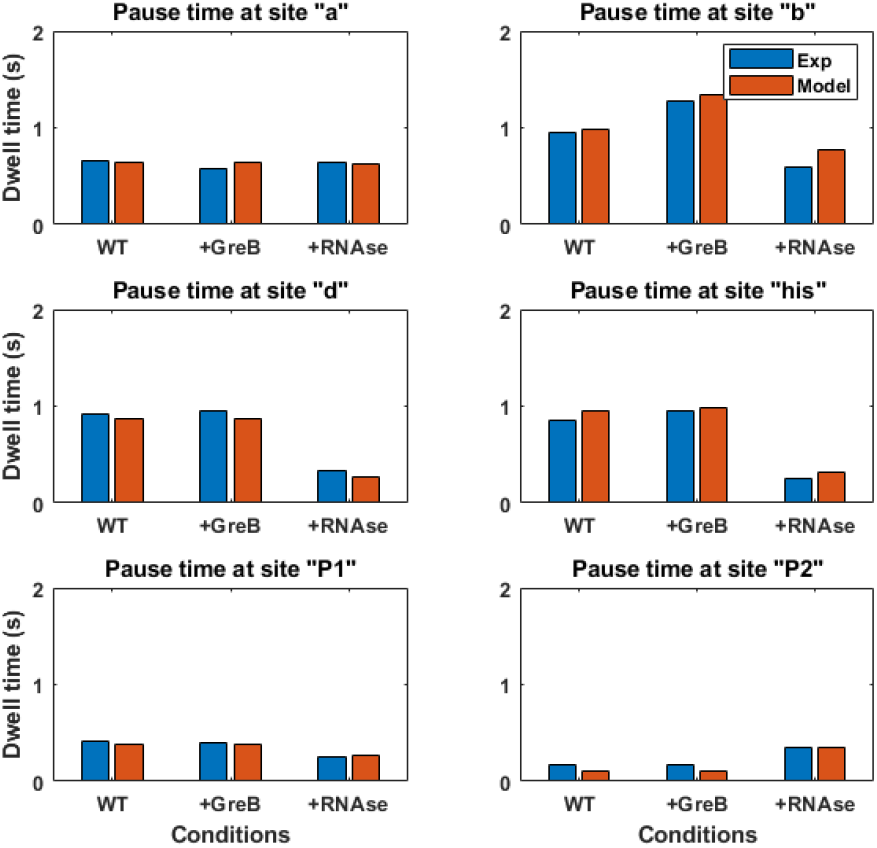
Averaged dwell times from experiments (blue) and model (red) at pause sites with various transcriptional factor conditions.

The effect of tension is modeled by introducing two different effective lengths *EL*_f_ and *EL*_bt_ for forward and backtracking translocation, respectively (Figure S6). Notice that the effective length for the forward translocation pathway is shorter than 1 base, while the external force acts on an effective length shorter than 0.1 base during backtracking (Table S2). The fitted values of effective length agree with previous work [13, 20]. These results indicate that opposing tension extends the averaged duration of backtracked pauses by decreasing the transcription rate and accentuating the entry into backtracked pausing. It also supports the idea that the entry into long-lived pauses, such as backtracked pauses, follows entry in short-lived pauses.

The predictive power of the model is demonstrated by the fact that it accurately predicts major pauses in the transcription of an unfamiliar 200 base sequence. This sequence preceding the repeat region of the 8XHis template was not included in the data used to optimize the model parameters. Figure 2c shows that the model successfully predicts the main pauses near bases 15, 45, 140 and 180 found experimentally by aligning transcription records and histogramming the dwell times.

To further test the validity of the model, we used Monte Carlo simulations to generate a large number of transcription traces, and we compared the dynamics of backtracking in experimental and simulated traces. The pauses at site ‘b’ in simulated traces were analyzed for backtrack depth and backtrack duration (Figure S7). The clear agreement between simulated and experimental results lends further support to the model.

## IV. STRENGTHS AND LIMITATIONS OF THE MODEL

The model identifies pausing sites and correctly characterizes the mechanism of transcription pausing. Note that, the model identifies sites of slow forward translocation rates as pre-translocation pauses. In other reports, these are often referred to as ubiquitous pauses. The model also distinguishes backtracked and hairpin-stabilized pauses, which are typically conflated as long-lived pauses.

Our results support a previous theoretical analysis of transcriptional pauses which suggests that a long-lived pause stabilizes from a short-lived ubiquitous pause [3]. For example, at pause site ‘b’, backtracking is favored over forward translocation because of the low forward translocation rate. Indeed, the energetic parameters of the model would predict comparable backtracking rates at the 35 bp site (pause ‘b’) and at the 190 bp site, but the fast forward translocation rate at the 190 bp site diminishes backtracking (Figure S3). Using the canonical Michaelis-Menten expression, we determined that the forward translocation rate along the template varies from less than 3 nt/s to 70 nt/s. This implies that a slowly transcribing complex may enter into a long-lived pause at one site, even if the back-tracking energy barrier at this position is higher than the barrier height at a position where transcription is faster.

The model predicts that the effective length of force is about one half of a base pair for the forward translocation pathway, but less than 0.1 base for the backtracking pathway. This result suggests that external forces insignificantly affect the backtracking rate. During backtracking RNAP must ratchet backwards on the DNA and disrupt the RNA-DNA hybrid near the active site. We hypothesize that the rate is determined in large measure by the denaturation of the hybrid complex. Thus, external forces cannot alter this process as much as biasing the equilibrium constant in the forward translocation pathway.

The hairpin-stabilized pause requires the interaction between a transcript hairpin and the RNAP flap domain. Previous models simulated the folding of nascent transcripts using the lowest-energy method [10, 13, 21]. However, that method may not locate the correct positions of hairpins, since RNA folds co-transcriptionally and may not readily reach the lowest-energy configuration for RNAP at the pause site. In addition, simulation of co-transcriptional RNA folding requires enormous computational resources, so we devised a new method, which considers the stability difference between a structure including a hairpin and the lowest-energy structure (*K_i_* in Figure 1d), to estimate the likelihood of the appearance of hairpin structures (SI). In this case, hairpins at position 101 and 178, although they are stable structures, are less likely to interact with RNAP than the less stable hairpins at positions 94 and 161, which correspond to pauses at sites ‘d’ and ‘*his*’, respectively (Figure S5). Our method also readily reproduced the pause ‘P2’, which is significantly lengthened in the presence of RNase.

The model can predict the transcriptional kinetics at pause sites ‘a’, ‘P1’, ‘b’, ‘P2’, ‘d’ and ‘*his*’. Nonetheless, the current model cannot characterize the pauses observed at site ‘c’ and at other less significant sites. The duration of the pause at site ‘c’ is largely unaffected by the addition of either GreB or RNase, suggesting a mechanism distinct from backtracking or hairpin-stabilized pausing that is not captured in the current model.

## V. COMPARISON WITH OTHER MODELS

The model described herein significantly extends earlier efforts to model the kinetics of transcription. Bai et al., Tadigotla et al. and others independently proposed models in which the kinetic of transcription is treated as a competition between the active transcription pathway and a branched pathway [9, 10, 12, 13, 21]. Although their models yield results in some statistical agreement with experimental results, the predicted pauses differ from those observed in single-molecule measurements. In addition, previous models mostly focused on short ubiquitous and Class II backtracked pauses, without considering Class I, hairpin-stabilized pauses and the effects of tension and transcription factors.

This new model treats the kinetics of transcription as a competition between one active and two branched pathways that differ significantly from previous efforts. In particular, backtracking is treated as a two-step mechanism in which the first step backwards must overcome a higher energy barrier than the successive steps. Unlike earlier models, the kinetics of Class I pauses are also included. By fitting specific kinetic parameters under specific experiment conditions, the resulting model achieves not only statistical agreement with experimental results, but reveals quantitative detail regarding the effects of DNA sequences, applied tension, and transcription factors.

## VI. CONCLUSION AND OUTLOOK

This purely thermodynamic consideration of the transcription complex accurately reproduces transcription kinetics. By incorporating both Class I and Class II pauses, the model refines our current understanding of active pathway and branched pathways in transcription and can be used to predict the occurrence of Class I and II pauses that regulate transcription.

Further improvements in our biochemical understanding of transcriptional pauses, in the quality of experimental data, and in the model itself, could improve the predictive power. For example, the model might predict the pause at site ‘c’ if the mechanism underlying this pause is determined and incorporated. Longer spans of high resolution transcription data would also improve optimization of the model and the accuracy of predictions by providing more sequence variations.

## ACKNOWLEDGMENTS

This work was supported by the National Institutes of Health (NIH) grants R01 GM084070 to LF. We are grateful to Carlos Bustamante and Alex Tong for generously providing high resolution transcription data.

