## Supplemental text and figures for "A thermodynamic model of bacterial transcription"

### Supplementary Information
### A thermodynamic model of bacterial transcription

Jin Qian,* David Dunlap,† and Laura Finzi‡
*Physics Department, Emory University*

#### I. A STATISTICAL APPROACH TO THE TRANSCRIPTION BUBBLE CONFIGURATION AND STATE TRANSITION

A transcription complex ($m$, $n$) is in a rapid equilibrium among many microstates, each defined by the parameter ($b$) which depends on the number of unpaired DNA bases upstream ($u$) and downstream ($d$) of the DNA-RNA hybrid inside the RNAP enzyme, the length of the hybrid ($h$) and the number of single-stranded RNA bases protected by RNAP ($r$) (Figure S1).

Equilibrium among microstates (dashed arrows in Figure S1) is reached rapidly compared to the time required for state transitions. Thus, for each transcription complex ($m$, $n$), the probability of a particular microstate $b$ is given by the Boltzmann distribution:

$$\mathrm{P_m^b} = \mathrm{Z_m^{-1}} \exp\left(\frac{-\mathrm{G_{TEC}^{m,b}}}{\mathrm{k_B T}}\right) \tag{S1}$$

$$\mathrm{Z_m} = \sum_{\mathrm{b}} \exp\left(\frac{-\mathrm{G_{TEC}^{m,b}}}{\mathrm{k_B T}}\right). \tag{S2}$$

The overall transition rate is calculated as

$$k_{\mathrm{m}\to\mathrm{m+1}} = \textstyle\sum_{\mathrm{b}} P_{\mathrm{m}}^{\mathrm{b}} k_{\mathrm{m}\to\mathrm{m+1}}^{\mathrm{b}} \tag{S3}$$

Figure S1 shows the forward translocation step as an example of statistical treatment in the model. Indeed, all the state transitions in the model are determined according to Equation (S3), as the summation of the products of the probability and the transition rate of individual microstates.

#### II. M-M EQUATION APPROACH FOR ACTIVE PATHWAY

The Michaelis-Menten equation (Equation (2) in the main article) is used to describe the forward translocation rate. The equation is derived from the Brownian-ratchet model, in which forward translocation occurs in three steps: (i) a fast equilibrium between position m and position m+1, (ii) recruitment of NTP at active site, (iii) catalysis and release of pyrophosphate. In this interpretation, $k_{\mathrm{max}}$ is the rate of NTP catalysis, $K_{\mathrm{d}}$ is the NTP

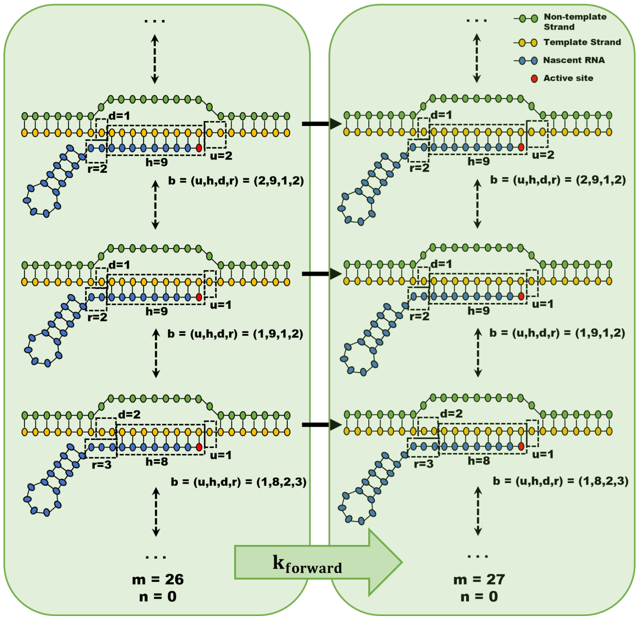


FIG. S1. An illustration of the statistical approach to transcription bubble configuration using the forward translocation step. Dashed arrows indicate fast equilibrium and solid arrows indicate the allowed state transitions.

dissociation constant, and $K_{\mathrm{i}}$ is the equilibrium constant between two adjacent translocation states. Yin and Steitz proposed another mechanism of forward translocation, in which the translocation is driven by the rotation of $O$ helix upon the release of pyrophosphate [1]. In this mechanism, the closed/open state of the $O$ helix determines the pre-/post-translocation state of the transcription complex. As shown in Figure S2, step 1 represents the incorporation of NTP with active site at position $m$; step 2 represents the transition from an open $O$ helix to a closed $O$ helix; step 3 represents the phosphoryl transfer reaction; step 4 represents the release of pyrophosphate, and step 5 represents the $O$ helix rotation which drives the translocation of TEC. Assuming the catalysis step 3 plus the PPi release step 4 are the rate-limiting steps, the rate of forward translocation is:

*
†
‡

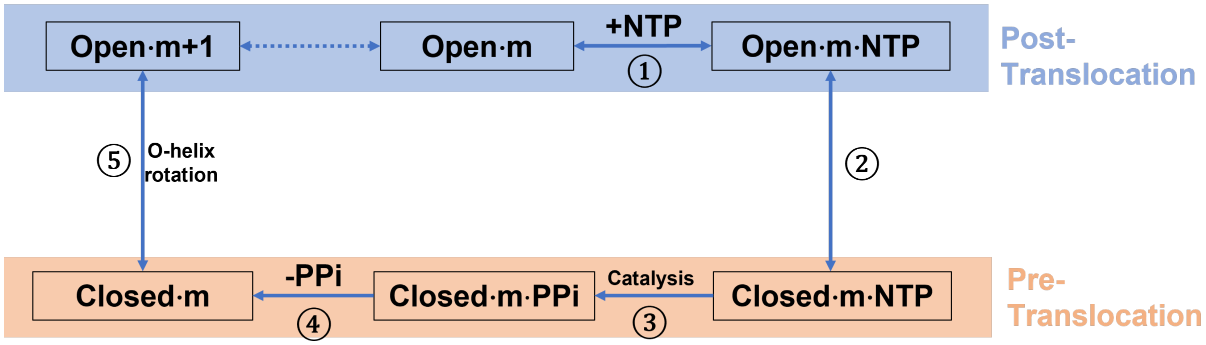


FIG. S2. A nucleotide addition cycle for an alternative mechanism proposed by Yin and Steitz, in which the translocation step is driven by the release of inorganic pyrophosphate.

$$k_{\text{forward}} = \frac{k_{\text{ppi-release}}\ k_{\text{catalysis}}[\text{NTP}]}{K_{\text{d}}\left(1 + exp\left(\frac{G_{\text{m+1}} - G_{\text{m}} + \Delta G_{\text{open}} - \Delta G_{\text{close}}}{k_{\text{B}T}}\right) exp\left(\frac{\Delta G_{close} - \Delta G_{open}}{k_{\text{B}}T}\right)\right) + [\text{NTP}]}$$
$$= \frac{k_{\text{max}}[\text{NTP}]}{K_{\text{d}}\left(1 + \ K_{\text{i}}\right) + [\text{NTP}]} \tag{S4}$$

In both cases, the M-M equation captures the mechanism of RNAP forward translocation, with the parameters $k_{\text{max}}$, $K_{\text{d}}$, and $K_{\text{i}}$ .

#### III. TREATMENT OF THE BACKTRACKING PATHWAY

As mentioned in the main text, the energy barrier for the first backtrack step is treated differently from subsequent backtrack steps, since it involves the disruption of the active site. We assume the energy barrier for an active TEC to enter the backtracked state to be:

$$\Delta \text{G}_{0\to-1} = \Delta \text{G}_{\text{bt}} - \text{G}_0 \tag{S5}$$

where $\Delta G_{bt}$ is a fixed activation energy specific for entering a backtracked state, $\Delta G_0$ is the energy of TEC at an active site. $\Delta G_{bt}$ is an energy barrier comparable to the free energy of base pairing lost during the formation of a transcription bubble. The rate constant to enter the backtracked state would be:

$$\text{k}_{0,\text{bt}} = \text{k}_1 \exp\left(-\Delta \text{G}_{0\to-1}/\text{k}_\text{B}\text{T}\right) \tag{S6}$$

where $\text{k}_1$ is the prefactor of backtracking. Equation (S6) indicates that backtracking is more likely to occur at a position with an unstable TEC bubble. However, the effect of TEC bubble energy may be compromised by the forward translocation rate at this position (Figure S3).

For any further backward translocation of RNAPs, the energy barrier should relate to the energy difference between two adjacent translocation states and the backtracked distance. Thus, for n > 0,

$$\Delta \text{G}_{-\text{n}\to-\text{n}-1} = \Delta \text{G}_{\text{bt\_increment}} + 0.5(\text{G}_{-\text{n}} - \text{G}_{-\text{n}-1}) \tag{S7}$$

and

$$\text{k}_{-\text{n},\text{bt}} = \text{k}_1 \exp(-\Delta \text{G}_{-\text{n}\to-\text{n}-1}/\text{k}_\text{B}\text{T}) \tag{S8}$$

where $\Delta \text{G}_{\text{bt\_increment}}$ represents the backtracking energy barrier due to increase in the length of the transcript inserted into the secondary channel.

Recovery from a backtracked state ($k_{btr}$) includes a diffusive and a cleavage pathway. The former occurs through RNAP diffusion, which also follows the Arrhenius equation with the energy barriers described above.

$$\text{k}_{-\text{n}-1,\text{btr}} = \text{k}_1 \exp(-\Delta \text{G}_{-\text{n}-1\to-\text{n}}/\text{k}_\text{B}\text{T}) \tag{S9}$$

and

$$\text{k}_{-1,\text{btr}} = \text{k}_1 \exp\left(-\Delta \text{G}_{-1\to 0}/\text{k}_\text{B}\text{T}\right) \tag{S10}$$

The cleavage pathway occurs by cleaving nascent RNA inserted into the secondary channel to register the 3' end of nascent RNA in the active site. This process is likely to be sequence-independent, and was assumed to occur at a constant rate.

#### IV. TREATMENT OF THE HAIRPIN-STABILIZED PAUSING PATHWAY

As discussed in the main text, the allosteric model suggests that a hairpin-stabilized pause is induced by the interaction between an RNA hairpin loop and the RNAP

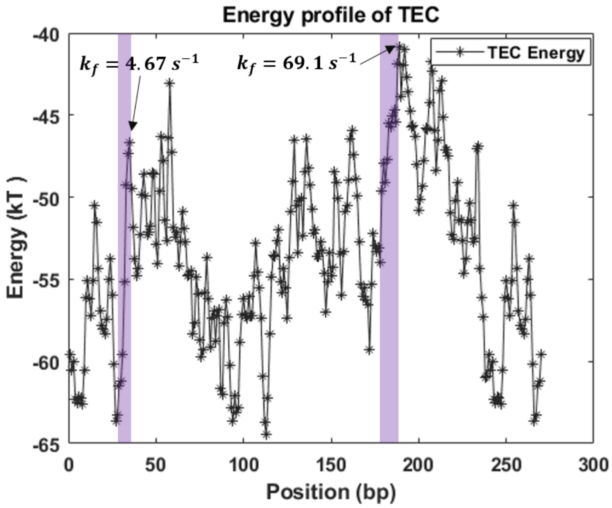


FIG. S3. Backtracking probability is highly dependent on the forward rate. The purple bands indicate two backtracking favored energy profiles. The one at 35 bp has a slow forward rate and causes the pause "b", while the one at 190 bp has a fast forward rate and shows no backtracking pause in both experiment and model fitting.

flap domain. We use Equation (S11) to model the entry rate to the hairpin-stabilized pause,

$$k_{hsp} = k_{on}/(1 + K_i) \tag{S11}$$

where $k_{on}$ is the catalytic rate of interaction between the RNA hairpin loop and the RNAP flap interaction, and $K_i$ is the fraction of hairpin helix formation.

The secondary structure of RNA transcript rapidly transitions among many microstates, and the simulation of transitions among these microstates is computationally expensive. We bypass this difficulty by simplifying the equilibrium to a two-state system of the lowest energy state and the hairpin-included state. The lowest energy state is determined by allowing all or at most a 100-nucleotide-long stretch of RNA outside of the exit channel to fold freely. A state including a hairpin is determined by first searching from the RNA 3' end for possible hairpin structures near the exit channel (up to 30 nt) before allowing all or at most 100 bases of the transcript to fold freely (Figure S4). The equilibrium between the lowest energy state and the hairpin state can be used to estimate the fraction of hairpin formation.

The model simulates the effect of RNase by shortening the length of freely folded RNA to 15 nt. The hairpin structures that induce pauses 'P1', 'd' and '*his*' disappear under this treatment, but a small hairpin structure, which is not normally favored, can readily form and induce pause 'P2'.

A chemical bond between the hairpin loop and the RNAP flap is required to stabilize the hairpin-flap interaction. The catalytic rate relates to the length of stem and loop, and the fraction of G and C in the loop as shown below,

$$k_{on} = k_1 \exp\left(-\frac{\mathrm{D}_{stem} * \Delta G_{stem} + \mathrm{D}_{loop} * \Delta G_{loop} + F_{GC} * \Delta G_{GC}}{kT}\right) \tag{S12}$$

where $k_1$ is the prefactor, $D_{stem}$ and $D_{loop}$ are the deviation from optimal lengths of stem (3 – 8 bases) and loop (4–20 bases), respectively, $F_{GC}$ is the fraction of G and C nucleotides within the loop, and $\Delta G_{stem}$, $\Delta G_{loop}$, and $\Delta G_{GC}$ are the energy changes due to $D_{stem}$, $D_{loop}$, and $F_{GC}$.

The exit rate from a hairpin-stabilized paused state ($k_{hspr}$) must be much slower than the entry rate, and is determined by the rate of RNAP hairpin denaturation. For simplicity, the rate is taken to be a constant.

#### V. SIMULATION OF DWELL TIME HISTOGRAM USING CONTINUOUS-TIME MARKOV CHAIN

With the transition rates between states, we can write the rate matrix of the Markov chain:

$$\mathbb{Q} = \begin{pmatrix} 1-\sum_{\mathrm{n}} k_{\{1,\mathrm{n}\}} & k_{0,\mathrm{bt}} & 0 & \dots & k_{\mathrm{hsp}} & k_{\mathrm{forward}} \\ k_{-1,\mathrm{btr}} + k_{\mathrm{cleavage}} & 1-\sum_{\mathrm{n}} k_{\{2,\mathrm{n}\}} & k_{-1,\mathrm{bt}} & \dots & 0 & 0 \\ k_{\mathrm{cleavage}} & k_{-2,\mathrm{btr}} & 1-\sum_{\mathrm{n}} k_{\{3,\mathrm{n}\}} & \dots & 0 & 0 \\ \vdots & \vdots & \vdots & \ddots & \vdots & \vdots \\ \vdots & \vdots & \vdots & \dots & \vdots & \vdots \\ k_{\mathrm{hspr}} & 0 & 0 & \dots & 1-\sum_{\mathrm{n}} k_{\{12,\mathrm{n}\}} & 0 \\ 0 & 0 & 0 & \dots & 0 & 1 \end{pmatrix} \tag{S13}$$

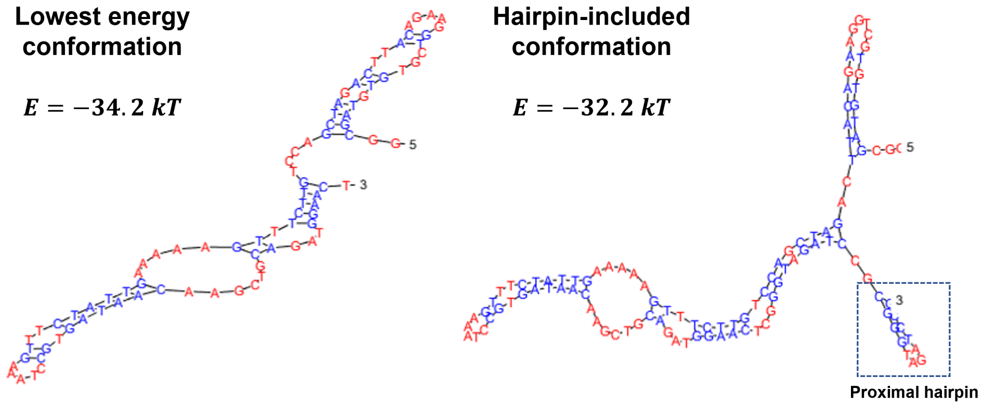


FIG. S4. Comparison of the lowest energy conformation with a conformation including a proximal (3') hairpin at position 12. Hairpin formation is unfavorable at this position.

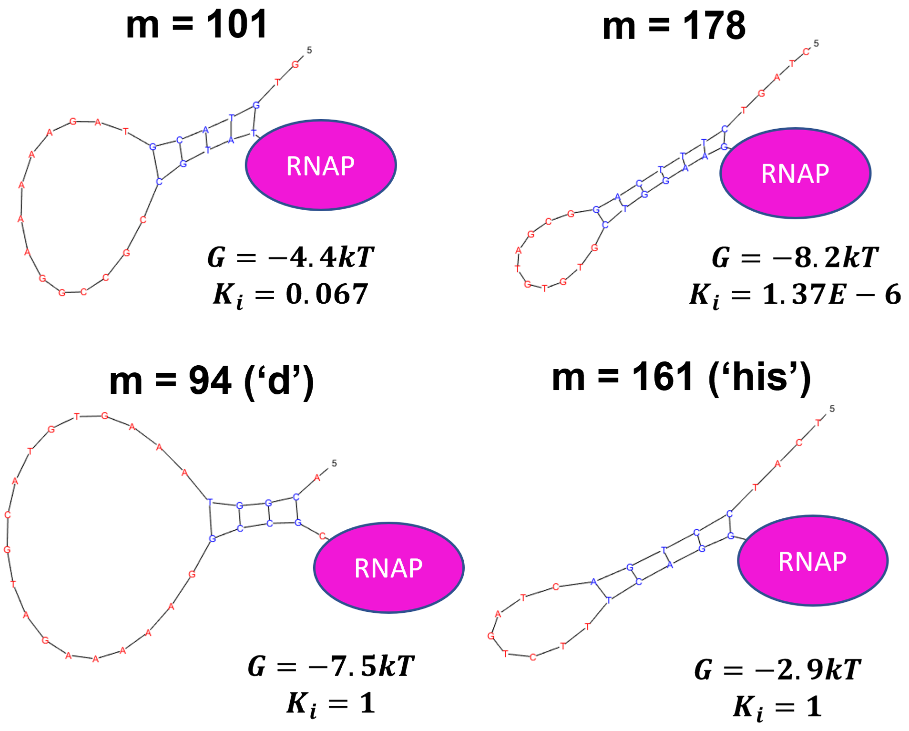


FIG. S5. Comparison of energy and $K_i$at different positions. Hairpin formation is unfavorable at position 101 and 178, although the hairpin structures at these positions are fairly stable. While at position 94 and 161, the hairpin structures can readily form and induce the hairpin-stabilized pauses.

The elements of row $n$ and column $m$ represent the transition rate from state $n$ to state $m$. The rows(columns) represent sequentially the active state, the backtracked states from 1 to 10 backtracking depth, the hairpin-stabilized state, and the next translocation state. The matrix is guaranteed non-singular. Given an initial state, which is clearly $[1, 0, 0, \ldots, 0, 0]$, the time spent in each

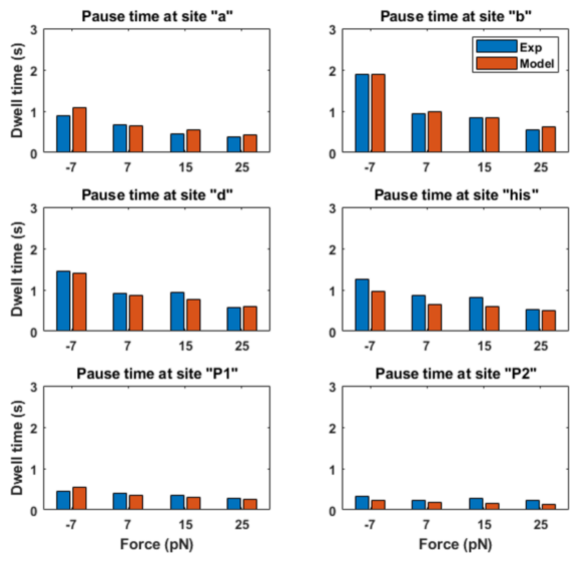


FIG. S6. Averaged dwell times from experiments (blue) and model (red) at pause sites with various forces conditions. The model accurately reproduced the pause times under all conditions.

state can be expressed as a matrix exponential

$$\begin{aligned}\overline{\tau} &= \alpha \exp\left(\mathrm{t}\mathbb{Q}\right) \\ &= \alpha \textstyle\sum_{n=0}^{\infty} \frac{t^n (VDV^{-1})^n}{n!} \\ &= \alpha V e^{Dt} V^{-1} \end{aligned} \qquad \text{(S14)}$$

where $\alpha$ is the initial distribution of states, $V$ consists of the eigenvectors of rate matrix $\mathbb{Q}$, and $D$ is a diagonal matrix of the diagonal elements of eigenvalues of $\mathbb{Q}$ ordered like the eigenvectors in $V$. Thus, the expected time spent in each state is

$$\overline{\mu} = \alpha V \lambda V^{-1} \qquad \text{(S15)}$$

where $\lambda$ is the negative inverse of the diagonal element of $D$ after replacing 0 eigenvalues with 1.

#### VI. OPTIMIZING THE MODEL WITH EXPERIMENTAL DATA

To optimize the model parameters, we first considered only the forward translocation pathway and fitted the equilibrium parameters $K_d$ , the kinetic parameters $k_{max}$ and an effective length $EL_f$ over which the external force acts during the forward translocation of RNAP. The result suggests that the pause at position ‘a’ is a pre-translocated pause which is consistent with the experimental data showing insensitivity to GreB. However, pauses at other sites were characteristically longer. In the next step, we included the backtracked pathway, the energy barriers $\Delta G_{bt}$, $\Delta G_{\mathrm{bt\_increment}}$ and an effective length $EL_{bt}$ for backtracking, maintaining the parameters of the forward translocation pathway set in the previous step (Table S2). The result suggests the backtracked pause is a large component of pauses at position ‘b’ but not at other positions, in agreement with the analysis of backtracking dynamics. Lastly, we included the hairpin-stabilized pause pathway with the rest of the parameters in the model fixed at the values identified in the preceding two steps (Table S2). The result gives a good agreement with pauses at other sites. We repeated the procedure above to fit the experimental data with GreB and RNase. Overall, the model faithfully reproduces pause times at all pause sites under all conditions with the exception of pause ‘c’ (Figure S6). Pauses at ‘c’ might originate from a different mechanism (see insights and limitations). Table S2 gives a list of the values of the fitted model parameters.

#### VII. MONTE CARLO SIMULATION AND ANALYSIS ON BACKTRACKING DYNAMICS

To further validate the model, we generated transcription data using Monte Carlo simulation and compared the dynamics of backtracking in the experimental and simulated traces. Figure S7a shows the example traces generated by using the optimized parameters shown in Table S2 under 10 pN of assisting tension. Only forward translocation and backtracking pathways were considered here for simplicity. The simulation was performed by calculating the probability of state transitions from the transition rates every 0.001 s. We collected the backtrack depth and duration from the simulated traces at pause site ‘b’ and compared them to the experimental results. Figure S7b and c shows that the simulated backtracking depth and duration was similar to that observed experimentally.

[1] Y. W. Yin and T. A. Steitz, The Structural Mechanism of Translocation and Helicase Activity in T7 RNA Polymerase, Cell **116**, 393 (2004).

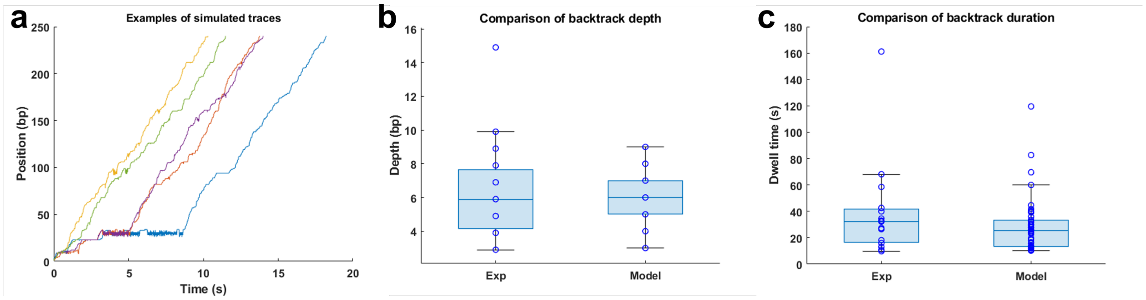


FIG. S7. Comparison between the backtracking dynamics in experimental and simulated data. (a) Examples of simulated traces generated by Monte Carlo simulation. (b) Distribution of backtrack depth observed experimentally and predicted by the model. (b) Distribution of backtrack duration observed in the experiments and predicted by the model.

TABLE S1. Summary of experimental pause positions and durations under different transcriptional factor conditions under 10pN and mechansims.

| Pause | Position of Peak (bp) | Averaged Duration (s) | | | Associated state(s) |
|---|---|---|---|---|---|
| | | WT | +GreB | +RNase | |
| 'a' | 9 | 0.66 | 0.58 | 0.64 | Pre-translocated |
| 'b' | 34 | 0.94 | 1.27 | 0.59 | Backtracked + Hairpin-stabilized |
| 'c' | 66 | 0.42 | 0.41 | 0.38 | Unknown |
| 'd' | 94 | 0.92 | 0.96 | 0.33 | Hairpin-stabilized |
| '*his*' | 161 | 0.85 | 0.95 | 0.25 | Hairpin-stabilized |
| 'P1' | 16 | 0.41 | 0.40 | 0.25 | Hairpin-stabilized |
| 'P2' | 44 | 0.16 | 0.17 | 0.34 | Hairpin-related, found to be hairpin-stabilized |

TABLE S2. Values (uncertainties) of the optimized parameters under 10pN assisting force and WT condition.

| Parameters and descriptions | | Symbol and Value | Note |
|---|---|---|---|
| Forward Translocation | Rate of NTP catalysis for AUCG | $k_{max} = [85(17), 77(15), 82(14), 41(7)]s^{-1}$ | Fitted |
| | Equilibrium constant for AUCG | $K_d = [34(7), 96(19), 15(4), 26(7)]\mu M$ | |
| | Effective length for forward translocation | $EL_f = 0.56(0.27)bp$ | |
| Backtracking | Prefactor of backtracking | $k_1 = 1000s^{-1}$ | Fixed |
| | Energy barrier height of first base-pair backtracking | $G_{bt} = 9.8(2.2)k_BT$ | Fitted with fixed $k_{max}$ and $K_d$ |
| | Energy barrier height of deeper backtracking | $G_{bt_{incre}} = 1.8(0.4)k_BT$ | |
| | Effective length for backtracking | $EL_{bt} = 0.06(0.04)bp$ | |
| Hairpin-stabilized pause | Energy change due to unlikely stem length | $\Delta G_{stem} = \text{Inf}$ | Fixed values |
| | Energy change due to unlikely loop size | $\Delta G_{loop} = \text{Inf}$ | |
| | Energy change due to GC fraction | $\Delta G_{GC} = 8.8(2.1)k_BT$ | Fitted with fixed $k_{max}$ and $K_d$ and backtrack related parameters |
| | Hairpin-flap interaction rate | $k_{on} = 807(292)s^{-1}$ | |
| | Hairpin denaturation rate | $k_{hspr} = 3.4(1.1)s^{-1}$ | |
| TEC structure | Allowed RNA-DNA hybrid length | $h = 7 \sim 9bp$ | Fixed range |
| | Allowed upstream spacer length | $u = 1 \sim 3bp$ | |
| | Allowed downstream spacer length | $d = 1 \sim 3bp$ | |
| | Allowed number of single-stranded RNA protected by RNAP | $r = 1 \sim 3bp$ | |
